# Attentional inhibition by alpha power is modulated by faster theta rhythm and audio-visual congruency during natural speech perception

**DOI:** 10.1101/2024.02.25.581860

**Authors:** Gabriel Byczynski, Hyojin Park

## Abstract

Audio-visual speech processing is a fundamental aspect of human communication; however, the neural oscillatory mechanisms, particularly those involving alpha rhythms, that underlie attention in tasks requiring attention and suppression remain unclear. To investigate this, we employed a complex audiovisual paradigm designed to explore how alpha rhythms, along with slower frequencies, monitor and integrate audio-visual information under congruent and incongruent conditions. Participants were presented with a TED Talk video while listening to auditory stimuli under four conditions: (1) congruent audio delivered to both ears, (2) congruent audio in one ear and incongruent audio in the other, with attention directed toward the congruent audio, (3) congruent audio in one ear and incongruent audio in the other, with attention directed toward the incongruent audio and (4) all-incongruent stimulus. To examine lateralized attention effects, participants were divided into left-attending and right-attending groups across individuals. By analysing fluctuations in alpha power with regards to audio-visual congruency and the side of attention, we observed a notable finding: alpha power fluctuations falling within the faster delta/theta range to the direction of attention, corresponding to task difficulty and accuracy. This result indicates a lateralized relationship between low-frequency rhythms and alpha-band activity, highlighting the role of alpha rhythms as a mediator of attention in audio-visual speech processing. Indeed, inhibitory alpha power fluctuation follows previous observations of sensory filtering. These findings underscore the importance of lateralized neural dynamics in tasks involving selective attention and suppression, providing new insights into the oscillatory mechanisms underlying audio-visual integration in human communication.

**Graphical Abstract:** 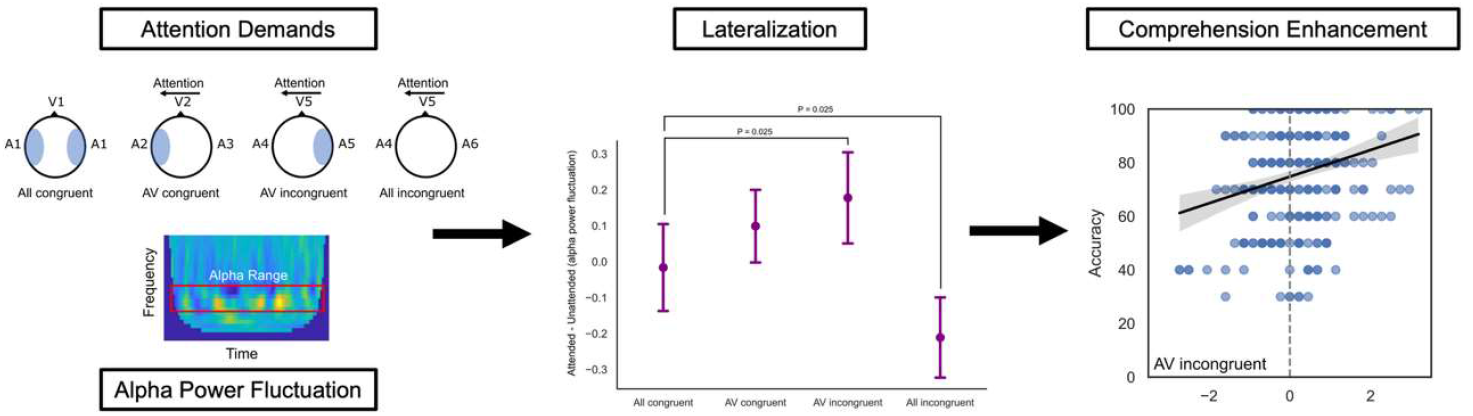

This work describes how alpha power fluctuation varies as a function of attention during complex audiovisual stimuli. We identified that alpha power fluctuation in the delta-theta range increase as a function of task demands and corresponds to comprehension accuracy. By understanding how temporal-spatial alpha power dynamics influence attention, we describe a novel cross-frequency attention mechanism.

## Introduction

Auditory and visual information are both important aspects of communication, and thus there is a wide field of research investigating how the brain perceives, processes, integrates, and stores auditory and visual information streams. To better understand audio-visual processing, and how the brain interprets both matching (congruent) and unmatching (incongruent) inputs between audio-visual stimuli, researchers utilize paradigms that require directed attention to specific modalities. One such investigation, known as the cocktail party paradigm, reproduces a real-life scenario in which auditory and visual information are provided, but not necessarily congruently (Golumbic et al., 2013; Park et al., 2016). This task produces scenarios in which either visual or auditory information is obscured or supplemented with other inputs, creating a complex scenario in which the brain must attend to a specific speaker. In these scenarios, several neural mechanisms become essential to extracting relevant information and suppressing the processing of irrelevant information. One such mechanism includes functional inhibition by alpha rhythm (Jensen & Mazaheri, 2010). During such functional inhibition, brain regions that are unrelated to the processing or storage of activity are inhibited, and thus information is actively directed through areas that remain engaged. Reasonably, paying attention to all incoming auditory and visual information in a given moment is metabolically and cognitively non-optimal, and thus attention can be considered as an act of resource management by directing relevant information to relevant brain areas (Oberauer, 2019).

There are numerous theories of how the brain directs attention, however here we focus on two major perspectives relevant to our hypotheses. The first suggests that attention enhances neural activity in areas responsible for processing stimuli. The second proposes that attention operates by suppressing activity in regions unrelated to the stimuli through inhibitory mechanisms (Foxe & Snyder, 2011). We expand on these hypotheses by considering that attention may not function as a continuous, steady-state process but rather as an intermittent, periodic sampling of the external environment. This sampling is suggested to follow rhythmic fluctuation (Fiebelkorn & Kastner, 2019; Helfrich et al., 2018). Fiebelkorn and Kastner (2019) proposed that theta oscillations (4–8 Hz) play a crucial role in visual attention by enhancing temporal resolution and facilitating periodic sampling of visual stimuli. Helfrich et al. (2018) also provided evidence that saccadic eye movements are synchronized with sensory input, and that this synchronization can be influenced by precisely timed stimulus presentations, indicating a rhythmic pattern in attentional sampling. While these findings have advanced our understanding of rhythmic attention in the visual domain, there is a lack of research exploring similar mechanisms in auditory processing. In this study, we investigate whether rhythmic attention operates in the auditory domain, focusing on low frequency (< 8 Hz) and its relationship with the role of alpha oscillations (8–12 Hz) in functional inhibition. We manipulate the lateralization of attention and the congruency between audio-visual stimuli to test our hypothesis.

Recent research has begun to address the gap in understanding rhythmic attention in the auditory domain. Plöchl et al. (2022) demonstrated that theta band activity (4–7 Hz) is associated with target detection in both visual and auditory modalities, providing evidence for theta oscillations in rhythmic attention. Additionally, alpha band activity appears to be coupled with theta rhythms in a counter-phasic manner, suggesting that alpha oscillations may inhibit task-irrelevant stimuli, potentially through theta-driven alpha modulation. In the visual domain, alpha rhythms are thought to reflect the sampling rate of perception and are cross-frequency coupled with theta during attention tasks (Samaha & Postle, 2015). Cross-frequency coupling (CFC) refers to the interaction between neuronal oscillations of different frequencies, where one frequency modulates another through various methods, including phase– amplitude, phase–phase, and amplitude–amplitude coupling (Jensen & Colgin, 2007). This phenomenon allows for the integration of information across different temporal and spatial scales in the brain, facilitating complex cognitive functions and enables the coordination of activity between distributed brain regions, serving as a mechanism for attentional control and the integration of sensory information (Canolty & Knight, 2010; Doesburg et al., 2012). Low-frequency oscillations, such as theta rhythms, have been linked to conditions requiring both auditory and visual monitoring, playing roles in attention and short-term memory (Keller et al., 2017). Therefore, theta-driven modulation of alpha activity may direct top-down attention by functionally inhibiting irrelevant information.

In our study, we employed a classical cocktail party paradigm to investigate audio-visual attention using magnetoencephalography (MEG). Our findings reveal asymmetric alpha power correlates of auditory attention within the theta frequency range. Consistent with previous visual domain studies, we observed that alpha band oscillations may serve as a ‘sampling rate’ mechanism, enhancing information intake. This mechanism appears to function by increasing sampling directed toward attended stimuli and/or decreasing sampling toward unattended or irrelevant stimuli. Further lateralization of this effect suggests regional contributions, underscoring the significance of top-down signaling through cross-frequency coupling in directing attention under complex audio-visual conditions.

## Materials and Methods

### Participants

We employed four experimental conditions as described in our previous study (Park et al., 2016). For complete methodology and detailed protocol, please refer to the original publication. Data were collected from 46 healthy right-handed participants (26 females, mean age 20.54 ± 2.58 years). All participants provided written consent and received monetary compensation for their participation. All participants had normal or corrected-to-normal vision and no history of developmental, psychological, or neurological disorders. Due to the nature of the task, only British English native speakers were recruited. Two subjects were removed, as one fell asleep and the other had excessive MEG noise. This left dataset from 44 participants (25 females, mean age: 20.45 ± 2.55 years). This study was approved by the local ethics committee (CSE01321; University of Glasgow, College of Science and Engineering) and conducted in conformity with the Declaration of Helsinki.

### Data acquisition

Neuromagnetic signals were obtained using a 248-magnetometer MEG system (MAGNES 3600 WH, 4-D Neuroimaging) in a shielded room with a sampling rate of 1017 Hz. Pre-processing included denoising using the Fieldtrip Toolbox (Oostenveld et al., 2011), and removal of electrooculogram (EOG) and electrocardiogram (ECG) artifacts using independent component analysis (ICA) and bad sensors according to the MEG analysis guidelines (Gross et al., 2013).

### Stimuli and task

Participants were presented with a visual stimulus and two auditory stimuli. The stimuli used were video and audio clips of a male speaker talking for between 7-9 minutes. Stimuli were sourced from TED talks (www.ted.com/talks/) and were edited to remove references to visual materials or referring to the gender of the speaker. Video clips were filmed by a professional filming company with a high-quality audio-visual device and recorded in 1920 x 1080 pixels at 25 fps (frame per second) for video and a sampling rate of 48 kHz for audio.

Three conditions were used in the present study: ‘All congruent’ condition, ‘AV congruent’ condition, ‘AV incongruent’ condition, and ‘All incongruent’ condition. Here, congruency refers to the alignment between auditory and visual stimuli. In the ‘All congruent’ condition, participants were presented with congruent audio-visual stimuli in both ears. ‘AV congruent’ and ‘AV incongruent’ conditions followed a dichotic listening paradigm, where participants received different auditory stimuli in each ear. In the ‘AV congruent’ condition, participants were presented with congruent audio-visual stimuli on either the left or right side and instructed to attend to the congruent audio. In contrast, the ‘AV incongruent’ condition maintained the same stimulus presentation as the ‘AV Congruent’ condition; however, participants were required to attend to the side where the audio was incongruent with the visual input (Figure 1A). Lastly, in the ‘All incongruent’ condition, participants were required to attend to a given side, while both visual and unattended audio were incongruent. Thus, participants in this final condition were presented with 3 streams of incongruent audivisual information. Participants were divided into two groups (22 participants each for the left- and right-attending groups), and a fixation cross color at the beginning of the visual stimulus cued them to attend to either the left or right auditory stimulus. Throughout the task, participants were instructed to maintain their gaze on a fixation cross, which was positioned over the speaker’s lips. Their eye movements and gaze behaviors were continuously monitored using an eye-tracking device to ensure compliance with the fixation requirement.

**Figure 1.**
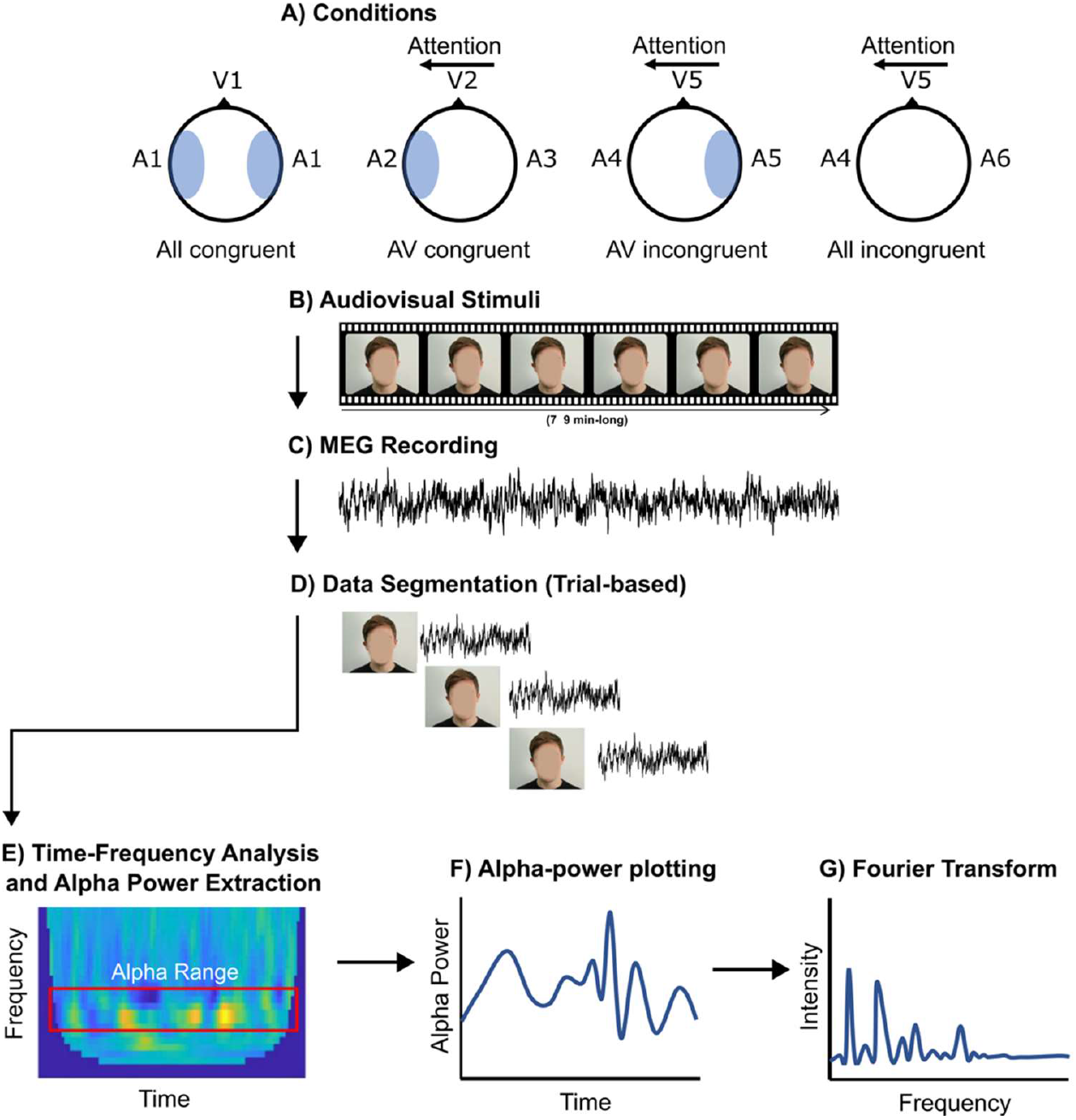
A) Conditions used in the present study, where the direction of attention is indicated by arrows, and the shaded region indicates the side in which auditory information is congruent with visual information (examples of the left-attending group). B) Audio-visual stimulus appearance. C) Continuous MEG recording during stimulus. D) MEG data were segmented into ∼3-second trials aligned with acoustically segmented speech chunks. E) Time-frequency analysis was performed for each trial. F) Alpha power was extracted over time, and G) Fast-Fourier Transform results on alpha power fluctuation.

After each condition, a questionnaire probing the content of the attended speech was delivered. We measured accuracy (%) of answers to measure comprehension of the attended stimulus.

### Extraction of alpha power fluctuation

We first acoustically segmented the speech recordings into discrete chunks using the Syllable Nuclei (de Jong & Wempe, 2009) script in Praat (Boersma & Weenink, 2018) (see Fig. 1D). Acoustic silences were identified based on a minimum pause duration of 0.25 seconds and a silence threshold of 25 dB. Each resulting speech chunk had a minimum duration of 1 second. The corresponding MEG data were then segmented to align with these speech chunks for subsequent analysis.

To determine the rhythms of the alpha power fluctuation in each condition, we used Fast Fourier Transform (FFT) of a subset frequency range of the alpha power. Using the FieldTrip toolbox (Oostenveld et al., 2011) in MATLAB2021b, data analysis was carried out following the Flux Pipeline for MEG analysis (Ferrante et al., 2022). Data were baseline corrected using -1.5 to 0 seconds before the onset of speech. After the time-frequency analysis, alpha power (8-12 Hz) was extracted from left and right auditory sensors in each participant. This produced a single trace representing the fluctuation of alpha power over time for the subset of auditory sensors. We then applied a Fast Fourier Transform (FFT) to determine at which frequencies the alpha power was oscillating (Figure 1). We investigated the peaks of FFT of alpha power fluctuation falling under theta and delta range (0.1 – 7 Hz).

### Statistical analysis

Statistical analyses were performed on data from all 44 participants, separated by attending direction (n = 22 per group). When comparing alpha power fluctuation, we used a repeated measures ANOVA (rmANOVA) with side of recording (left or right auditory hemisphere) as the repeated measure, and group (left or right attention) and peak number as the between-subject factor. We calculated the difference between attended and unattended-side alpha power fluctuation with simple subtraction within-subject for ANOVA analysis across conditions. To understand the relationship between alpha power fluctuation frequency and behaviour, we lastly analyzed the effect of condition and alpha power fluctuation on comprehension accuracy. We used a mixed effects regression model with accuracy as function of the difference in alpha power fluctuation between attended and unattended sides, condition, and peak as fixed effects, and random intercepts for subjects.

For statistical differences between the attending groups shown in Figure 5, cluster-based permutation tests were performed in Fieldtrip (Oostenveld et al., 2011). An independent samples t-test was calculated for each sensor-time sample with an alpha threshold of 0.05 as the cluster-building threshold (two-tailed). Only significant results (*p* < 0.05) are reported.

## Results

### Alpha power fluctuation is faster in the right-hemisphere during All congruent conditions

We first analyzed the All congruent condition, in which participants were presented with congruent audiovisual information on both sides. Importantly, while participants were not asked to attend to a given side at this point, the analysis considered the subsequent group assignment to investigate pre-existing differences between groups in terms of attention. There was no effect of hemisphere on alpha power fluctuation (F(1,294) = 0.099, *p* = 0.753), and no main effect of group (direction of attention) (F(1,294) = 3.180, *p* = 0.076), thus there was no pre-existing alpha power fluctiation tendency toward the side of attention in the subsequent conditions (Figure 2A). There was however a significant interaction between hemisphere and group (F(1,294) = 13.441, *p* < 0.001). Post-hoc testing with Bonferroni correction which aimed to determine within-group differences revealed that both groups had greater alpha-power fluctuation to the right side compared to the left (Right attending group: t(294) = 2.370, *p*_*corr*_ = 0.036; Left attending group: t(294) = 2.815, *p*_*corr*_ = 0.01) (Figure 2B). This indicated that in conditions where participants are not asked to attend to a specific side, there was consistently greater alpha power fluctuation to the right, possibly indicating basal attentional preferences under free task demands, which is consistent with previous reports of right-lateral bias during auditory attention (Schmithorst et al., 2013).

**Figure 2.**
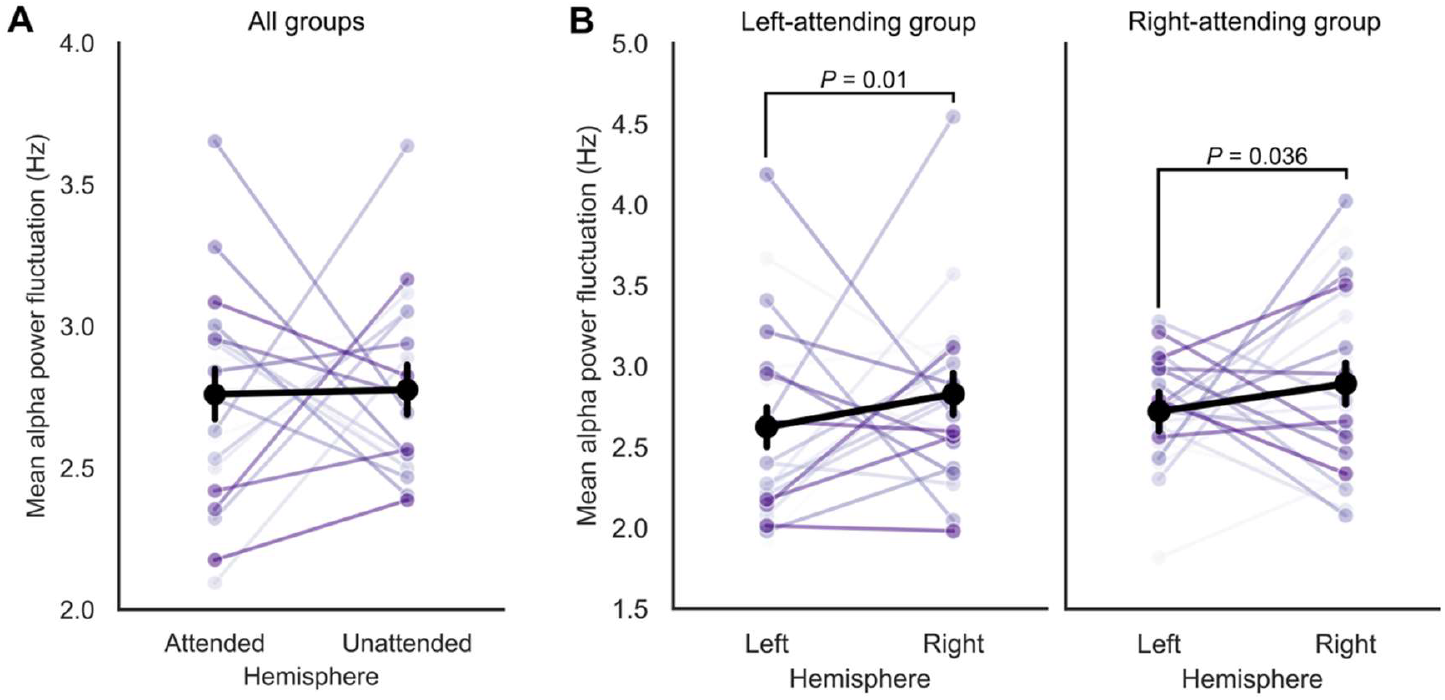
A) Overall effect of hemisphere on mean alpha power fluctuation across both groups by future attention direction. B) Alpha power fluctuation by group and hemisphere during All congruent task conditions by hemisphere. Errorbars denote standard error. Individual datapoints show mean fluctuation frequency. Aggregated means by subject are shown.

### Faster alpha power fluctuation in the attended side during the AV congruent and incongruent conditions

We next investigated how directing attention influences alpha power fluctuation in each hemisphere. During the AV congruent condition, participants are asked to attend to one side in which the delivered audio is congruent with the visual stimulus. To the unattended side, participants are delivered incongruent audio. We found a significant effect of hemisphere on alpha power fluctuation, (F(1,294) = 4.632, *p* = 0.032) (Figure 3A), whereby the attended hemisphere had faster alpha power fluctuation than the unattended hemisphere (mean difference = 0.099 Hz, SE = 0.046 Hz, t(294) = 2.14, *p* = 0.032). There was no interaction between group and hemisphere (F(1,294) = 1.820, *p* = 0.178), indicating that this effect was not specific to a given hemisphere. There was however, an effect of group (F(1,294) = 12.308, *p* < 0.001), in which the left-attending group had lower alpha power fluctuation overall than the right-attending group (mean difference = 0.179 Hz, SE = 0.051 Hz, t(294) = 3.508, *p* < 0.001).

**Figure 3.**
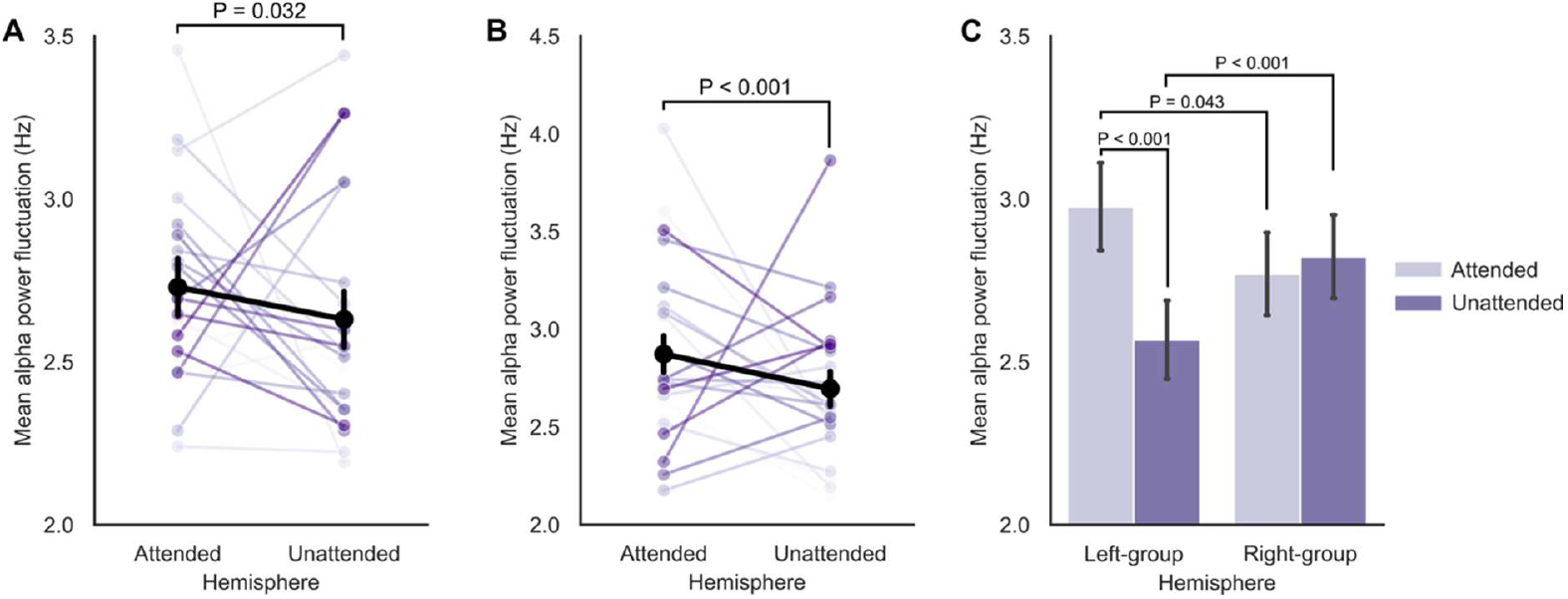
A) AV congruent condition alpha power fluctuation by side of attention. B) AV incongruent condition alpha power fluctuation by side of attention. C) AV incongruent condition interaction between group and hemisphere. Individual datapoints are shown for pointplots. Errorbars denote standard error. Aggregated means by subject are shown.

During the AV incongruent condition, participants were asked to attend to audio on a given side, and were presented with visual stimulus which was congruent with the *unattended* auditory stimulus. This condition was therefore more difficult, as it required processing of auditory information with conflicting visual information, as compared to the AV congruent condition. There was a main effect of hemisphere (F(1,294) = 11.426, *p* < 0.001) (Figure 3B), and a significant interaction between group and hemisphere (F(1,294) = 19.273, *p* < 0.001) (Figure 3C). There was no main effect of group, indicating that overall, alpha power fluctuation did not differ between direction of attention. Post-hoc testing revealed again that the attended side showed faster alpha power fluctuation (mean difference = 0.178 Hz, SE = 0.053 Hz, t(294) = 3.380, *p* < 0.001). For the group-hemisphere interaction, there was a significant difference between the left attended and right attended sides (mean difference = 0.207, SE = 0.084, t(294) = 2.461, *p* = 0.043), a significant difference between the left attended and right unattended side (Left attending group) (mean difference = 0.409, SE = 0.074, t(294) = 5.495, *p* < 0.001), and a significant difference between the left and right unattended sides (mean difference = 0.255, SE = 0.067, t(294) = 3.800, *p* < 0.001). This indicated that in the more challenging task conditions, participants indicated faster alpha power fluctuation to the side of attention, and further, that the left group boasted strong differences between the attended and unattended side compared to right-attending participants.

### Faster alpha power fluctuation in the unattended side during the All incongruent condition

Lastly, we analyzed the All incongruent condition, in which all visual and auditory stimuli were incongruent. Thus, participants were presented with 3 distinct stimuli. We found only a significant effect of hemisphere (F(1,294) = 18.389, *p* < 0.001), however, unexpectedly, in this condition there was faster alpha power fluctuation to the incongruent side (mean difference = 0.211, SE = 0.049, t(294) = 4.288, *p* < 0.001).

### Differences in alpha power fluctuation are driven by condition difficulty and correspond to accuracy

We analyzed the difference in alpha-power fluctuation between the attended and unattended sides across all conditions. We found a significant effect of condition on difference in fluctuation (F(3,882) = 11.533, *p* < 0.001), in which post-hoc testing revealed that the AV incongruent condition and All incongruent condition had significantly different alpha power fluctuation imbalance between hemispheres compared to the All congruent (control) condition. Specifically, the AV incongruent condition indicated more attending-lateralized alpha power fluctuation (t(294) = 2.688, *p*_corr_ = 0.025), while the All incongruent condition boasted more unattending-lateralized alpha power fluctuation (t(294) = 2.748, *p*_corr_ = 0.025) (Figure 4A).

**Figure 4.**
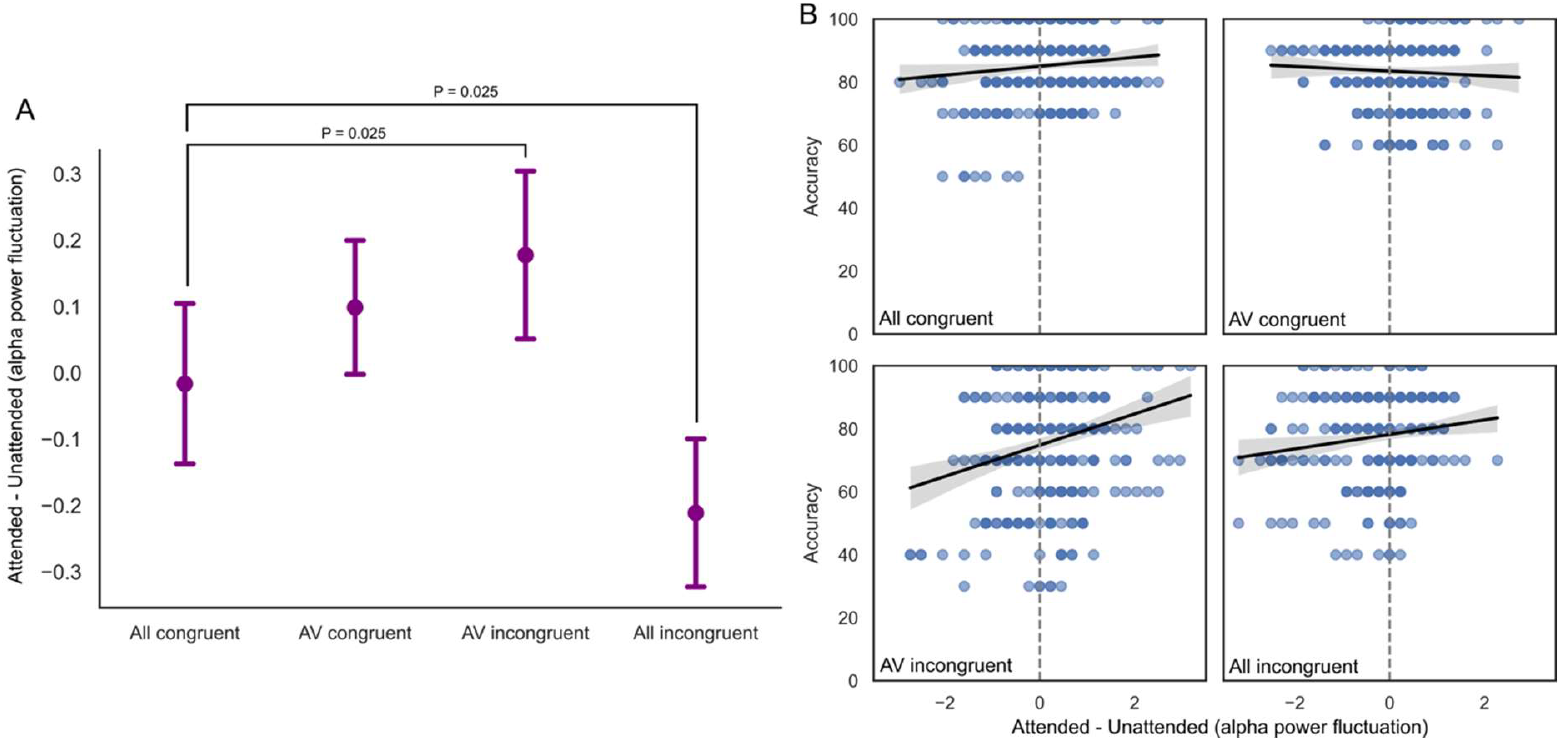
A) Difference in alpha power fluctuation between attended and unattended hemisphere across 4 conditions. Errorbars indicate standard error. B) Relationship between alpha power fluctuation frequency differences between hemispheres, and comprehension accuracy. Shaded area indicates 95% CI.

**Figure 5.**
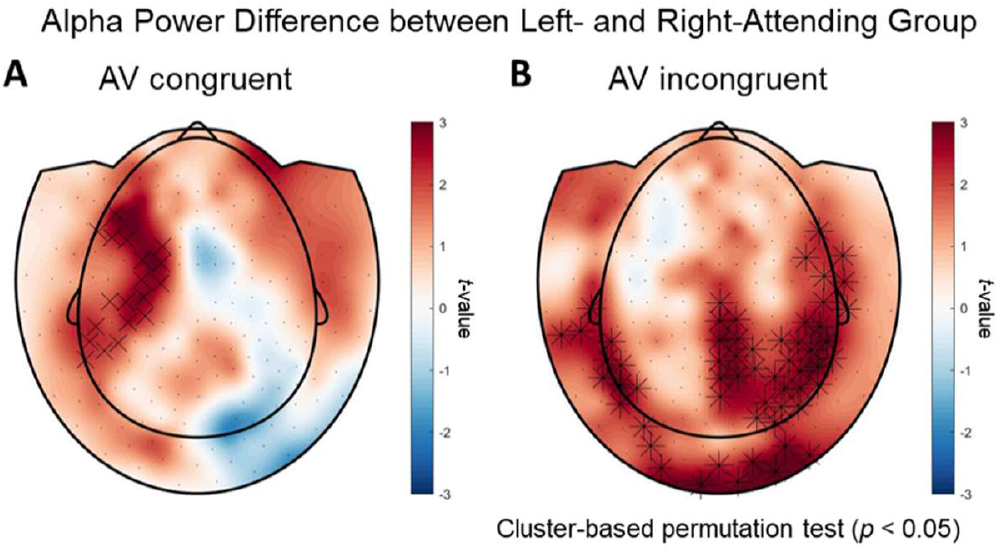
Topographical maps of alpha power (8–12 Hz) differences between the left-attending and right-attending groups for each condition: AV congruent (A) and AV incongruent (B). Statistical comparisons between the groups were performed separately for each condition using two-tailed cluster-based permutation tests (*p* < 0.05). Significant effects were observed between 2-2.5 seconds after the onset of the speech chunks for both conditions.

Lastly, we sought to understand the relationship between alpha power fluctuation and comprehension across conditions, in order to confirm if the observed pattern of alpha power fluctuation frequency increase is relevant to ecological comprehension. Using a mixed effects regression, we found the overall model significantly improved fit compared to a null model (χ^2^(8) = 145.72, p < 0.0001). There was a significant effect of condition on comprehension-accuracy-change from the all congruent condition, whereby the AV incongruent condition signficantly reduced accuracy by 6.9% (SE = 1.04%, *p* < 0.001), and the All incongruent condition by 9.98% (SE = 1.03%, *p* < 0.001). The AV congruent condition did not produce a significant change in accuracy (*p* = 0.14) from the All congruent condition. There was a significant interaction between alpha power fluctuation and the AV incongruent condition (β = 2.30, SE = 1.15, *p* = 0.045), indicating evidence of a relationship between alpha power fluctuation and accuracy occuring only during the AV incongruent condition (no other condition had a significant interaction, all *p* > 0.05) (Figure 4B).

### Global alpha power differences between attending groups across the sensor array

The difference between the attending groups prompted further investigation of global alpha (8-12 Hz) power differences between the left- and right-attending groups for each audiovisual conditions: AV congruent (Figure 5A) and AV incongruent (Figure 5B). Statistical comparisons between the groups were conducted separately for each condition using cluster-based permutation tests (*p* < 0.05), applied at 500 ms time intervals. For the AV congruent condition, alpha power was significantly increased in the left auditory and temporal areas in the left-attending group compared to the right-attending group, suggesting enhanced suppression of the to-be-ignored speech on the right side. For the AV incongruent condition, alpha power was significantly increased in the central and bilateral parieto-occipital regions, as well as in the auditory and temporal areas, suggesting suppression of lip movements that were congruent with the to-be-ignored speech. Significant effects were observed between 2-2.5 seconds after the onset of the speech chunks for both conditions.

## Discussion

In this study, we investigated how alpha-band (8–12 Hz) oscillations are modulated by attention and audio-visual congruency during natural speech perception. By analyzing alpha power fluctuations and overall alpha power under congruent and incongruent listening conditions, we aimed to understand the role of oscillatory dynamics - particularly cross-frequency interactions - in supporting selective attention in complex audio-visual environments.

Our findings reveal two key aspects of alpha dynamics: first, alpha power fluctuation differed between the attended and unattended hemispheres for all except the All congruent condition. Specifically, the attended hemisphere on average displayed faster alpha power fluctuation in the delta-theta range. These results align with previous theories proposing that attentional sampling may operate rhythmically, modulated by low-frequency oscillations such as theta (Fiebelkorn & Kastner, 2019; Helfrich et al., 2018). It may represent an increased temporal sampling of perceptual information which becomes necessary when visual information no longer provides congruent information. This pattern has been observed before, wherein sensing organs alternate between operational modes resulting in low and high modal frequencies (Morillon et al., 2019), and in the visual domain as well (Samaha & Postle, 2015). Second, overall alpha power also differed between attending groups. In the AV congruent condition, alpha power was significantly greater in the left auditory and temporal areas of the left-attending group compared to the right-attending group, supporting the notion that alpha oscillations facilitate functional inhibition of task-irrelevant inputs (Foxe & Snyder, 2011; Jensen & Mazaheri, 2010). Interestingly, in the AV incongruent condition, alpha power increases were observed not only in auditory areas but also in bilateral parieto-occipital regions, suggesting that suppression was extended to visual inputs - particularly lip movements congruent with the to-be-ignored auditory stream. Such findings support broader models of cross-modal suppression during audiovisual processing (Janssens et al., 2018; Ro, 2019), and illustrate how alpha-band activity is recruited flexibly depending on the sensory modality and attentional demands. As such, alpha dynamics subserve distinct cognitive mechanisms: fluctuation frequency may govern temporal sampling of attended information (Morillon et al., 2019; Samaha & Postle, 2015), while overall alpha power may regulate inhibition of distractors (Jensen & Mazaheri, 2010). These two dimensions of alpha control may operate independently depending on task complexity.

Moreover, the faster alpha fluctuation rate falling into the delta-theta range raises the possibility of cross-frequency coupling between theta phase and alpha amplitude, a phenomenon proposed to coordinate information processing across temporal scales (Canolty & Knight, 2010; Doesburg et al., 2012; Jensen & Colgin, 2007). Phase-amplitude coupling between delta/theta oscillations and alpha power has previously been shown in other studies (e.g., Cohen et al. (2009)), and is proposed to represent a coordinated timing of neuronal firing within local networks (Szczepanski et al., 2014). Taking this effect into consideration, our findings align with those of Gomez-Ramirez et al. (2011), suggesting that delta-entrained rhythms can modulate alpha power during auditory and visual paradigms. While our study does not directly investigate phase-amplitude coupling but rather examines the frequency of alpha power, it is possible that the observed alpha power oscillation results from delta/theta-phase modulated alpha-amplitude coupling.

Interestingly, alpha power fluctuation also differed as a function of condition, with more demanding conditions (see accuracy reporting in Park et al. (2016)) resulting in greater attended vs unattended alpha power fluctuation differences. We also found in the AV incongruent condition, which boasted the greatest difference between attended and unattended hemisphere alpha power fluctuation frequency, the degree of difference positively correlated with measures of comprehension accuracy, tying together the neurobiological evidence for attentional upregulation with perceptual differences in comprehension. Indeed this suggests a meaningful effect of alpha power fluctuation on semantic information processing. The only exception to this effect was the all incongruent condition, in which participants were exposed to three streams of incongruent visual and auditory information. Here, we speculate that other mechanisms for both visual and auditory suppression compensate, however the inversion of faster alpha power fluctuation to the unattended side requires further investigation.

Our results suggest that the speed at which the alpha power oscillates (perhaps via operationalizing theta/delta-oscillation frequency), reflects a top-down attentional mechanism of information processing that could reflect the temporal sampling theory in the visual domain. Faster theta-modulated coupling results in faster information sampling, thus preferentially directing and attentional control of incoming information.

Lastly, the non-signficant difference of significant alpha fluctuation changes under the AV congruent condition compared to the All congruent condition suggests that top-down attentional modulation may be less necessary when auditory and visual streams are congruent and mutually supportive. When audiovisual inputs align, attentional demands may decrease, and inhibition through alpha dynamics may not be actively recruited (Klatt et al., 2020; Park et al., 2016). This interpretation is supported by our previous behavioral findings showing that comprehension performance is higher under congruent audiovisual conditions compared to incongruent ones (Park et al., 2016). This also explains why the AV incongruent condition evoked the strongest difference, as it could reflect greater attentional resource recruitment. Furthermore, the AV incongruent condition requires the supression of congruent auditory and visual information, while the All incongruent condition does not present any audiovisual congruency. It could be in this difference, that the reversal of alpha power fluctuation rate can be explained, as the condition requires no supression of congruent audiovisual information, that is, no suppressed audiovisual integration. This also may suggest that the mechanism of attentional sampling is specific to audiovisual integration, however this should be investigated further.

In summary, our results demonstrate that alpha oscillations play a flexible, dynamic role in supporting selective attention during complex audiovisual speech perception. Faster fluctuations of alpha power reflect enhanced temporal sampling when attentional control is necessary, while increases in overall alpha power indicate suppression of irrelevant information across sensory modalities. These findings extend current models of attention by highlighting how cross-frequency dynamics and hemispheric asymmetries contribute to the neural basis of communication in naturalistic environments.

## Conclusion

Our results provide the first evidence that alpha power fluctuation frequency is modulated by delta/theta rhythm when greater attentional control is required during a naturalistic speech perception task. Our study suggests a novel neural cross-oscillatory correlate of complex auditory attention in association with audio-visual congruency. Future research may further investigate specifically the directionality of delta/theta coupling with alpha power fluctuation and explore the plausibility of this fluctuation as an information ‘filter’ which appears to be lateralized.

## Acknowledgments

We would like to thank Joachim Gross and Ole Jensen for their helpful and engaging discussions that contributed to this study.

